# Fast anther dehiscence state recognition system establishing by deep learning to screen heat tolerant cotton

**DOI:** 10.1101/2021.11.09.467902

**Authors:** Zhihao Tan, Jiawei Shi, Rongjie Lv, Qingyuan Li, Jing Yang, Yizan Ma, Yanlong Li, Yuanlong Wu, Rui Zhang, Huanhuan Ma, Yawei Li, Li Zhu, Jie Kong, Xianlong Zhang, Wanneng Yang, Ling Min

## Abstract

Cotton is one of the most economically important crops in the world. The fertility of male reproductive organs is a key determinant of cotton yield. The anther dehiscence or indehiscence directly determine the probability of fertilization in cotton. Thus, the rapid and accurate identification of cotton anther dehiscence status is important for judging anther growth status and promoting genetic breeding research. The development of computer vision technology and the advent of big data have prompted the application of deep learning techniques to agricultural phenotype research. Therefore, two deep learning models (Faster R-CNN and YOLOv5) were proposed to detect the number and dehiscence status of anthers. The single-stage model based on YOLOv5 has higher recognition efficiency and the ability to deploy to the mobile end. Breeding researchers can apply this model to terminals to achieve a more intuitive understanding of cotton anther dehiscence status. Moreover, three improvement strategies of Faster R-CNN model were proposed, the improved model has higher detection accuracy than YOLOv5 model. In addition, the percentage of dehiscent anther of randomly selected 30 cotton varieties were observed from cotton population under normal temperature and high temperature (HT) conditions through the integrated Faster R-CNN model and manual observation. The result showed HT varying decreased the percentage of dehiscent anther in different cotton lines, consistent with the manual method. Thus, this system can help us to rapid and accurate identification of HT-tolerant cotton.

**One sentence summary:** The deep learning technique was applied to identify the anther dehiscence state for the first time to quickly screen heat tolerant cotton varieties and help to explore key genetic improvement genes.

## Introduction

Cotton is an economically important crop, and its reproductive development is susceptible to a variety of adverse stresses that affect its yield and quality. The reproductive organs of cotton include stamens and pistils, and stamens are more sensitive to heat stress than female organs (Peet et al., 1998). In many summer crops, reproductive organ abortion caused by high temperatures is manifested by normal development of the female reproductive system and abnormal development of the male reproductive system, failure to produce functional pollen or failure of the anthers to achieve dehiscence properly to release pollen. Anther development is a complex process, going from sporogenic cells to anther dehiscence, and has been divided into 14 periods by studying a variety of male sterile mutants (Sanders et al., 1999). Anther dehiscence, the final step in anther development, includes three processes: secondary thickening of the inner wall of the anther chamber, degradation of the septum cells, and dehiscence of the cleft which ultimately allow the release of pollen (Kim et al., 2010). Therefore, anther dehiscence is directly related to the probability of fertilization in cotton. If we can obtain phenotypic data on anther dehiscence quickly and accurately and conduct genome-wide association analysis, we can easily obtain the functional genes related to the anther dehiscence. It is also important to analysis the molecular mechanism of cotton male reproductive organ respond to stresses.

In the past, the acquisition of cotton dehiscent or indehiscent anther number data from the pictures relied mainly on visual observation and manual counting, it is difficult to guarantee the accuracy of visual readings because anther growth is intermingled, resulting in unclear definition of individual anthers, and the background and foreground of anthers are easily confused. Moreover, a larger amount of anther data is needed to judge the anther growth and dehiscence status of individual plants in population under different conditions. However, it is obviously difficult to achieve this accurately and quickly with manual methods.

With the development of computer vision technology and plant phenome platforms, machine learning-based image processing techniques are widely used. However, in the training process of machine learning, there is a need to manually extract image features and feed the obtained classification features into the classifier for learning after a weighting process. Due to the poor generalization ability of classifiers and the need for large amounts of supporting data, the shortcomings of machine learning methods have gradually been exposed in the process of agricultural intelligence development.

In 2012, the concept of deep learning was proposed, and deep learning techniques have evolved rapidly in the past few years. Image recognition techniques based on deep learning and convolutional neural networks are gradually replacing machine learning-based image processing techniques in a wide range of fields. Through classification and extraction of image features and end-to-end training of deep learning models, computers can accurately detect specific content in images. Through the building of different datasets and the replacement of deep learning network architectures, researchers can obtain network models that are more suitable for research purposes than previous approaches.

In this study, using YOLOv5 (Redmon et al., 2016; Redmon and Farhadi, 2017; Redmon and Farhadi, 2018; Pang et al., 2020; Xu et al., 2020) and Faster R-CNN (Ren et al., 2017) models, combined with a variety of data augmentation methods, a cotton anther recognition model based on deep learning was obtained. This model can quickly batch collected cotton anther images for recognition, detect the dehiscent and indehiscent anthers, and obtain their phenotypic data.

Before the advent of deep learning, the usual machine target recognition process required human preprocessing of images before target detection and included cropping, augmentation, and segmentation. Various features of the image were extracted and handed over to a support vector machine (SVM) classifier (Piccialli and Sciandrone, 2018) for learning and detection. However, manual preprocessing is time-consuming and labor-intensive, and after the features are extracted, feature screening and evaluation must be performed according to the actual situation, and the weights of various features in the learning model must be artificially adjusted to achieve the best recognition effect. Against the background of the current state of machine intelligence, the disadvantages of traditional machine learning are obvious; the preliminary work requires considerable manual labour, the recognition accuracy is not ideal, and the technique is difficult to use in actual production.

After 2012, image recognition techniques based on deep learning and convolutional neural networks gradually replaced machine learning-based image processing techniques in a wide range of fields. The YOLO series, Faster-RCNN and single shot multibox detector (SSD) (Liu et al., 2016) are three important deep learning neural network models. Faster-RCNN mainly crawls preselected boxes and then performs deep learning classification. The image detection process of Faster-RCNN includes crawling region proposal, candidate feature frame extraction, and candidate feature frame classification. First, the convolution data of whole image is obtained. Then, the data is automatically fed into a region proposal network (RPN) (Zhu et al., 2021) to obtain the features of candidate regions. Finally, the features are classified by a softmax classifier and then adjusting for some special classes using a regressor. Faster-RCNN is a big improvement over its previous two generations: Fast-RCNN and RCNN (Girshick et al., 2014) in terms of recognition accuracy and speed. The CNN family of deep learning models is one of the mainstream models and has demonstrated powerful functionality in many fields, such as image detection and semantic segmentation. However, the YOLO model more cleverly uses the idea of regression by taking the whole image as input, dividing it into several boxed regions, removing individual boxes with very low relevance by setting specific thresholds, and finally selecting the highest scoring region with a nonmaximum suppression algorithm. The model removes the boxes that overlap with it until all alternative boxes are traversed, yielding the final output. In addition, many scholars are studying lightweight network structures. For example, MobileNet (Howard et al., 2017) and SqueezeNet are applied to YOLO networks to further improve their speed of detection and create the possibility of transplanting YOLO models to portable devices to ensure the accuracy of recognition as much as possible.

To date, no reports of machine learning-based anther identification systems in academia, but the application of target detection technology to agriculture using machine learning has been very extensive (Barre et al., 2017; Fuentes et al., 2017; Ubbens and Stavness, 2017; Gutierrez et al., 2019), which gives us great incentive to build a deep learning-based anther identification system for cotton. In maize, a parabolic model has been used to mine the diversity of stem-end meristematic tissues and to find candidate genes that correlate with the transport of phytohormones, cell division, and cell size by GWAS (Yang et al., 2007). In rice, the ratio of spikes to leaves, a new trait of rice, has been extracted using a feature pyramid network mask model that has achieved leaf and spike recognition accuracies of 0.98 and 0.99, respectively (Yang et al., 2020). Ferentinos KP has designed a convolutional neural network model to solve the problem of early plant disease detection. Through the deep learning method, several model structures have been trained with plant leaf images and have identified the corresponding plant leaf lesions with 99.53% accuracy. The model has become a powerful tool for the early diagnosis and early warning of plant leaf diseases and can be further improved. Therefore, the system can be used in real time in a real cultivation environment (Ferentinos, 2018). Ubbens JR et al. have designed an open source deep learning tool called Deep Plant Phenomics for plant phenotypic deep learning. This tool provides pretrained neural networks for several common plant phenotypic tasks including leaf counting, image classification and age regression. Botanists can use the neural networks provided and trained by this platform to train their plant phenotypes (Ubbens and Stavness, 2017). Nikita Genze et al. have proposed a convolutional neural network-based seed germination status recognition system that can automatically identify seed categories (including maize, rye, and pearl millet) in petri dishes and automatically determine whether the seeds are germinating. The system achieves an average accuracy of 94% on test data and can help seed researchers to better determine seed quality and performance (Genze et al., 2020). Scientists use hyperspectral imaging technology to collect spectral and image information from maize seeds and combine convolutional neural networks and support vector machines to model and train spectral data sets and image data sets. This model can quickly detect the vigor state of seeds and simultaneously predict their germination status, providing a framework to advance research on seed germination (Pang et al., 2020). A MobileNetv2-YOLOv3-based model that combines pretraining methods such as hybrid training and migration learning to improve its generalization for the early identification of tomato leaf spot disease has been proposed (Liu and Wang, 2021). Image processing and machine learning techniques have been used to accurately classify the three stages of plant growth and soil for different germplasms of two species of red clover and alfalfa. The accuracy on test data was shown to be more than 90% (Samiei et al., 2020).

In addition to their applications in computer vision, deep convolutional neural networks can be used in agricultural production, and they have broad application prospects in the combination of natural language processing (NLP) and agriculture (Schmidhuber, 2015). Protein ubiquitylation is an essential posttranslational modification process that plays a critical role in a wide range of biological functions. Siraj et al. proposed a method of predicting plant ubiquitin sites by using a hybrid deep learning model with a deep learning neural network and long-term and short-term memory. This method uses protein sequences and physical and chemical properties as the input to the model and approximate ubiquitin sites as its output. In ten cross-validations, the highest accuracies were shown to be 81% and 82%. This method improves the current situation of wasted time and person power when using traditional experimental methods to predict plant ubiquitin sites (Siraj et al., 2021). J Wekesa et al. propose a multi-feature fusion prediction model based on deep learning that combines categorical boosting and extra trees into a single meta-learner. The model is used to predict the function of plant long noncoding RNAs (lncRNAs). Experiments on *Zea mays* and *Arabidopsis thaliana* have yielded 0.9820 and 0.9652 areas under precision/recall curves (AUPRCs), respectively (Wekesa et al., 2020). OA Montesinos-López have implemented a multi-trait deep learning model with a feed-forward network topology and a rectified linear unit activation function with a grid search approach for the selection of hyperparameters. This model covers the multi-trait prediction of grain yield, days to heading and plant height. The results indicate that the deep learning method is a practical approach for predicting univariate and multivariate traits in the context of genomic selection (Montesinos-López et al., 2019).

## Results

### YOLOv5 model design

YOLOv5 is a typical one-stage detection model, which increases the detection speed by 50% compared with the previous generation YOLOv4, and its model size is only 1/10 of that of the previous generation model. The adaptive anchor frame calculation and the use of Focus structure enhance the accuracy of the model for small target recognition. At the same time, the model has four network models with different depths, and the best balance between detection accuracy and recognition speed can be found. It is very common for cotton anthers to block each other in the image, and the obscured anthers are easily ignored in the final output of the prediction box. YOLOv5 uses the soft-NMS (Bodla et al., 2017) method when screening the prediction box.

Since cotton anthers overlap and obscure each other, the use of the NMS algorithm results in an inability to accurately identify adjacent anthers, and only the anther images with the highest confidence are retained. Therefore, we use the soft-NMS algorithm. The idea of the NMS algorithm is that for a certain category X having N candidate boxes, the candidate boxes are sorted by their confidence, and the highest confidence box A is selected. The other candidate boxes Bi (i=1, 2, 3…) are compared with the highest confidence box A, and an IoU threshold is set. If its IoU is higher than this threshold, the candidate box B1 is discarded. Then the candidate box B2’s IoU is compared with that of the highest confidence box A. After several iterations, only prediction boxes that have an IoU lower than the set IoU value are retained. Although this method can prevent the same target from being repeatedly selected by multiple prediction boxes, it cannot prevent overlapping or occluded targets from being ignored.

The idea of Soft-NMS is that M is the current highest scoring box and Bi is the pending box. The larger the IoU of Bi and M, the more the score Si of Bi drops, rather than having the score go directly to zero as in NMS. This method can effectively retain anther images that overlap and ensure the accuracy of identification results. The linear weighting formula for Soft-NMS can be expressed as:

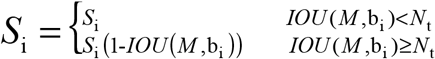

### Faster R-CNN model design

Faster R-CNN is a classical two-stage object detection network. The network model structure is mainly composed of four parts: feature extraction, region proposal, classification, roi pooling and its comprehensive performance has been greatly improved, especially for the detection accuracy of small targets. The cotton anther belongs to the range of small target detection in the whole image, so we trained the Faster R-CNN model to identify the anther dehiscence state has a better detection effect.

Conv layers is a classical CNN network target detection method, mainly includes three layers of conv, pooling, relu, usually uses to extract the feature maps of the input image. The extracted feature maps will be called by subsequent region proposal networks and classification networks. In the convlayers structure, contains 13 conv layers, 13 relu layers, 4 pooling layers. The Faster R-CNN has a very ingenious detail in the convlayers, it does augmentation treatment on all convolutional layers, fills a layer in the outer layer of the input matrix, so that the matrix is larger than it was, and the images that have been treated in this way are deconvoluted again, and after the convolution operation, the image is kept consistent with the size of the input image. The matrix size is unchanged when the image goes through the conv layer and relu layer, and will change to 1/2 of the original size after going through the pooling layer, so that when going through the convlayers structure, the size of the input matrix changes to 1/16 of the original size, so that the resulting feature maps can all correspond one-to-one with the original graph.

Conventional detection methods usually use a sliding window or the selective search method to acquire detection frames, whereas Faster R-CNN discards traditional methods and directly generates detection frames using region proposal networks, which greatly enhances the detection frame generation speed. The region proposal network structure is actually divided into two processes, the first process by softmax classification anchors, to obtain foreground and background (detection target is foreground), the second process is used to calculate the bounding box regression offset for anchors to obtain the exact proposal. Finally, the proposal layer is responsible for integrating foreground anchors and bounding box regression offset to obtain proposals, while simultaneously removing proposals too small and beyond the boundary. The entire Faster R-CNN network arrives to proposal layer, completing detection targets, the next two structures are mainly image recognition.

For the traditional CNN network, the input image of the model must be fixed size, and the output of the model must be a fixed vector or matrix. In practical applications, there are two solutions for images of different sizes: cut the picture to a fixed size or warp the image to a fixed size. However, these solutions will either cause the loss of image information, or lead to changes in the shape information of the image. Therefore, the structure roi pooling is proposed in Faster R-CNN to solve the problem of different image size. Roi pooling is mainly responsible for collecting feature maps and proposal boxes, calculating proposal feature maps, and sending it to the subsequent identification layer. First of all, proposal is mapped to the same scale as feature maps, and then the vertical and horizontal directions of each proposal are divided into seven parts, so that the output of different sizes of proposal is 7*7, realizing fixed-length output.

Classification using the obtained proposal feature maps, the structure calculates which category each proposal belongs to through full connect layers and softmax, and outputs the probability vector; at the same time, the position offset of each proposal is obtained again by bounding box regression, which is used to return a more accurate target detection box.

The loss function of the object detection network of Faster R-CNN is shown in Formula:

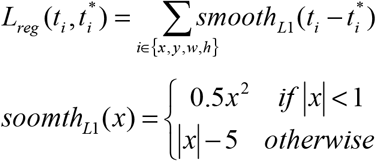

In the above-mentioned formula, i represents anchors index; t represents predict bounding box; t* represents ground true box corresponding to positive anchor; x, y, w, h represents center point coordinates of box, width, height.

### Data augmentation

#### Auto augment

This approach creates a search space for data-enhanced policies in which a policy contains many sub-policies and randomly selects one sub-policy for each image in a small batch data set. Each sub strategy consists of two operations, each of which is an image processing function similar to flatting, rotation, or shearing, and the probability and magnitude of applying those functions, using a search algorithm directly on the data set to find the best data augmentation strategy.

#### Random Resize

Random Resize scales the new image to the same pixel size as the original image by randomly clipping the original image in the data set according to the random aspect ratio.

#### Random Flip

Random Flip is a common way of data augmentation, which generates new data set samples by randomly flipping the original image of the data set up and down or left and right.

#### Mixup

Mixup is a data augmentation method for mixing two samples and label data at their corresponding ratios and then generating new sample and label data. Suppose x_1_ is a sample of batch one, y_1_ is the label corresponding to the sample of batch one; x_2_ is the sample of bach two, y_2_ is the sample corresponding label of bach tow, x_mix_ and y_mix_ is the newly generated sample and corresponding label, respectively. λ is the mixing coefficient resulting from the hyperparametric α and β conducted beta distributions. The principal formula of the mixup method can be expressed as:

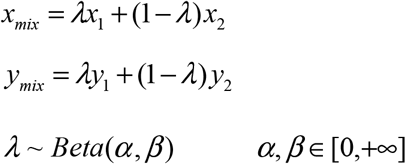

According to the study, we know that as the hyperparameters α and β increase, the error and generalization ability of the network training will increase. When the beta distribution of the mixing coefficient λ is α=β=0, the network recovers to the ERM (Empirical Risk Minimization) principle to minimize the training data average error; the beta distribution of the mixing coefficient λ has the best generalization ability and robustness. This method can make full use of all the pixel information, but at the same time also introduces some unnecessary pseudo-pixel information.

#### Cutmix

Cutmix (Yun et al., 2019) is by cut some regions in the sample and randomly fill in the pixel values of other samples in the data set, and at the same time distribute the final classification results according to a certain proportion. Compared with mixup, cutmix can prevent the occurrence of non-pixel information in the training process. Filling the pixel information of other regions with the missing area of cut can further enhance the positioning ability of the model. At the same time, this method will not increase the training and reasoning burden of the model.

#### GridMask

By generating a mask with the same resolution as the original image, GridMask multiplies the mask with the original image to get a new image. The pixel value of the new image in the fixed area is 0, which is essentially a regularization method. Compared with directly changing the network structure, GridMask only needs to be augmented when the image is input.

#### Normalized

We usually use this method after data augmentation, normalizing the pixel value of the image and scaling the pixel value to [0,1] can prevent the attributes of the large value interval from excessively dominating the attributes of the decimal value interval, and at the same time avoid the numerical complexity in the process of calculation.

In this study, the data augmentation process was shown in the Figure 1.

**Figure 1:**
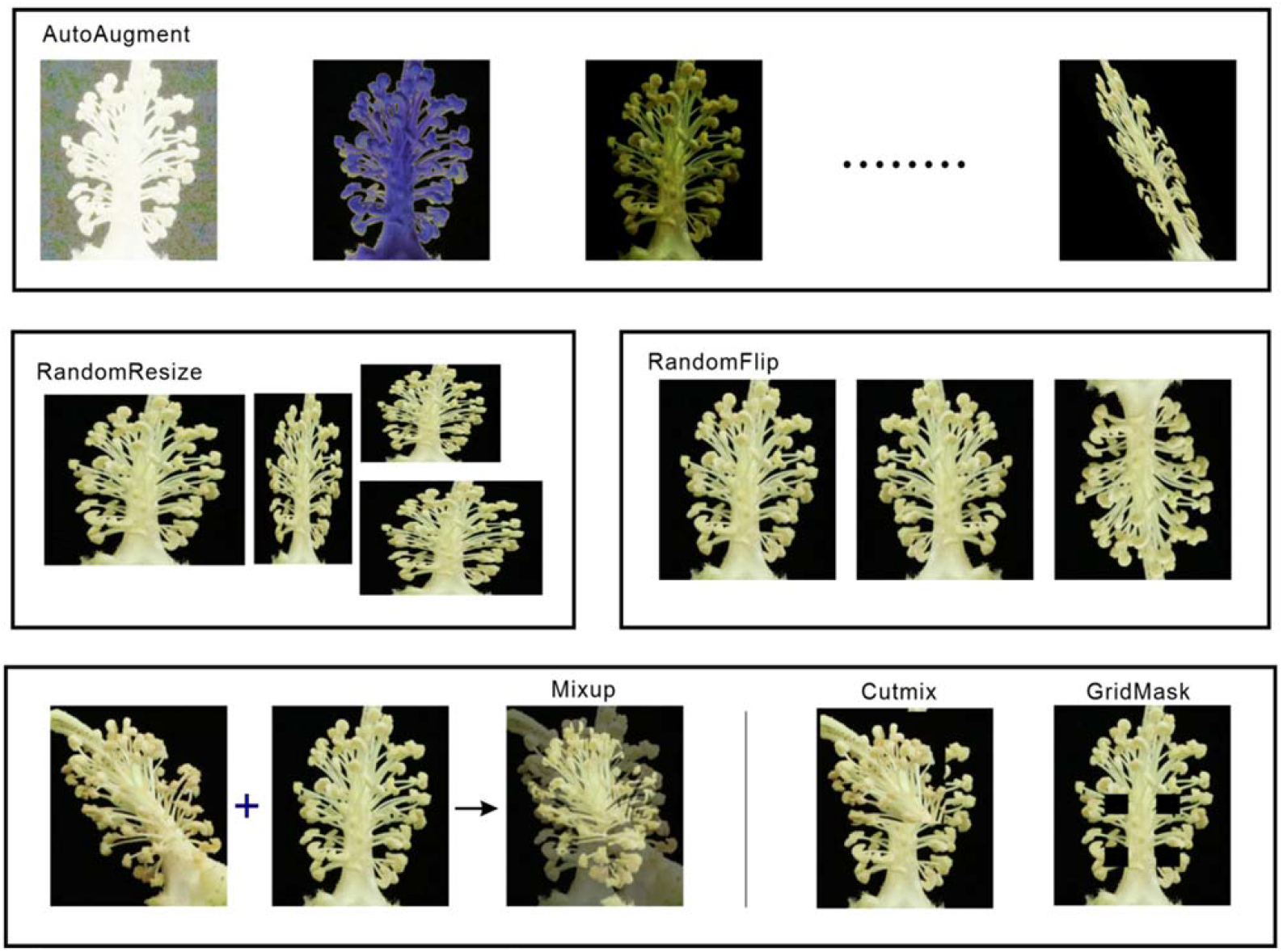
Data augmentation. The images above showed the effect of the same cotton anther image processed with different data augmentation methods.

### Model training

In this study, comparative experiments and control variables were used, YOLOv5 and Faster-RCNN models were used, and various data demonstration methods such as mixing and mixed cutting were used to train for sample imbalance, so as to verify the performance of different models and training methods on the same evaluation index validation set. Firstly, the self-made data set was segmented and analyzed, and VOC format was used to store training set, test set and verification set. Secondly, the model was trained according to whether the data demonstration algorithm was added. Finally, cosine strategy was used to periodically attenuate the learning rate. The training ends when the average loss remains stable. In this study, the training process of Faster R-CNN model was shown in the Figure 2.

**Figure 2:**
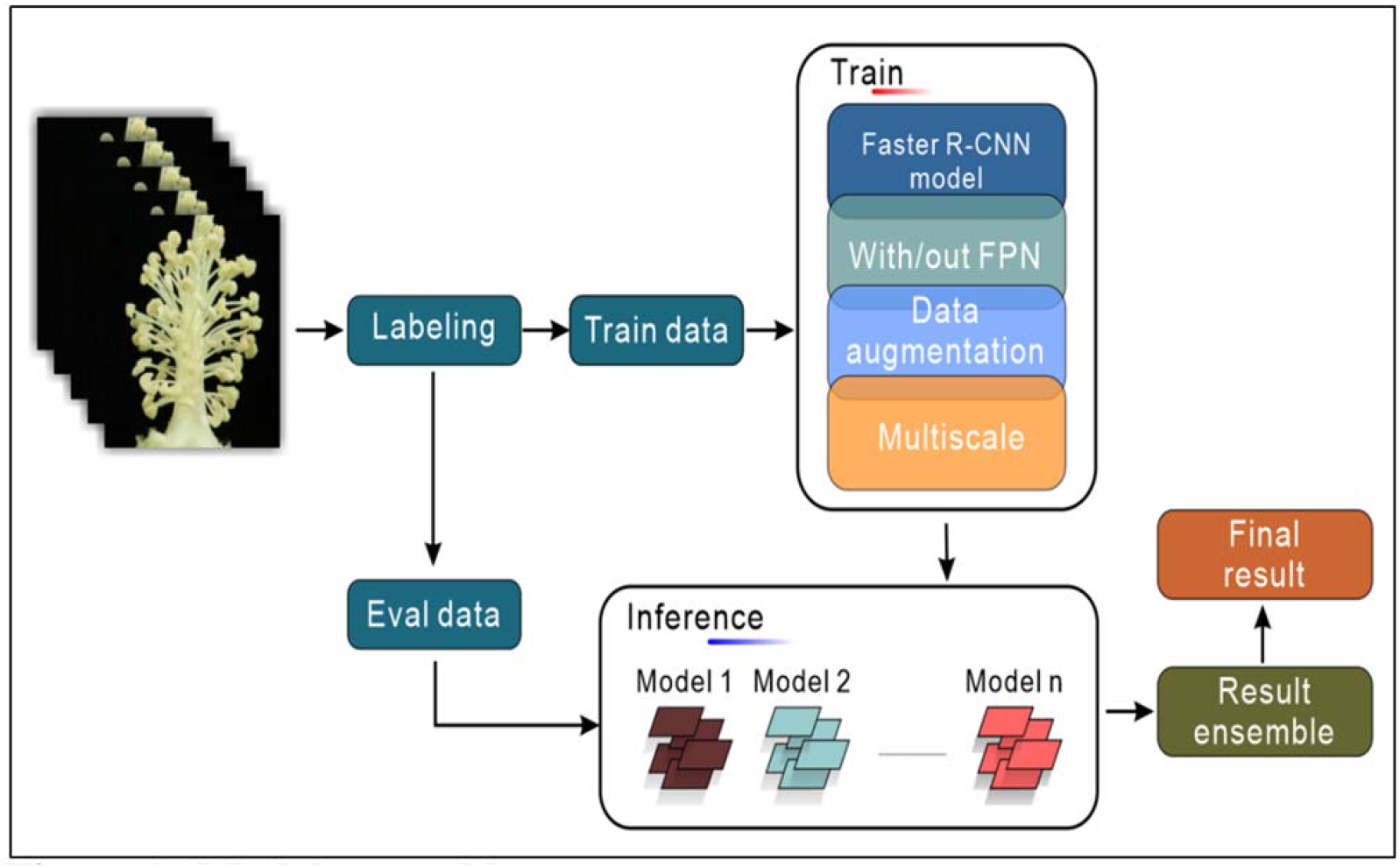
Model ensembles. Integrated flow chart of cotton anther recognition model ensembles.

The models obtained by different training strategies were tested on the test set and the prediction results of multiple models were obtained. The results of the four groups of comparison experiments indicated that the proposed Faster R-CNN neural network with data augmentation and FPN structure and Multi-Scale could effectively detect the dehiscence and indehiscence in cotton anther images. Compared with other methods, this method has significant advantages in recognition accuracy. The recognition effect was shown in the Figure 3. The final result was obtained by the prediction results ensembles of multiple models.

**Figure 3:**
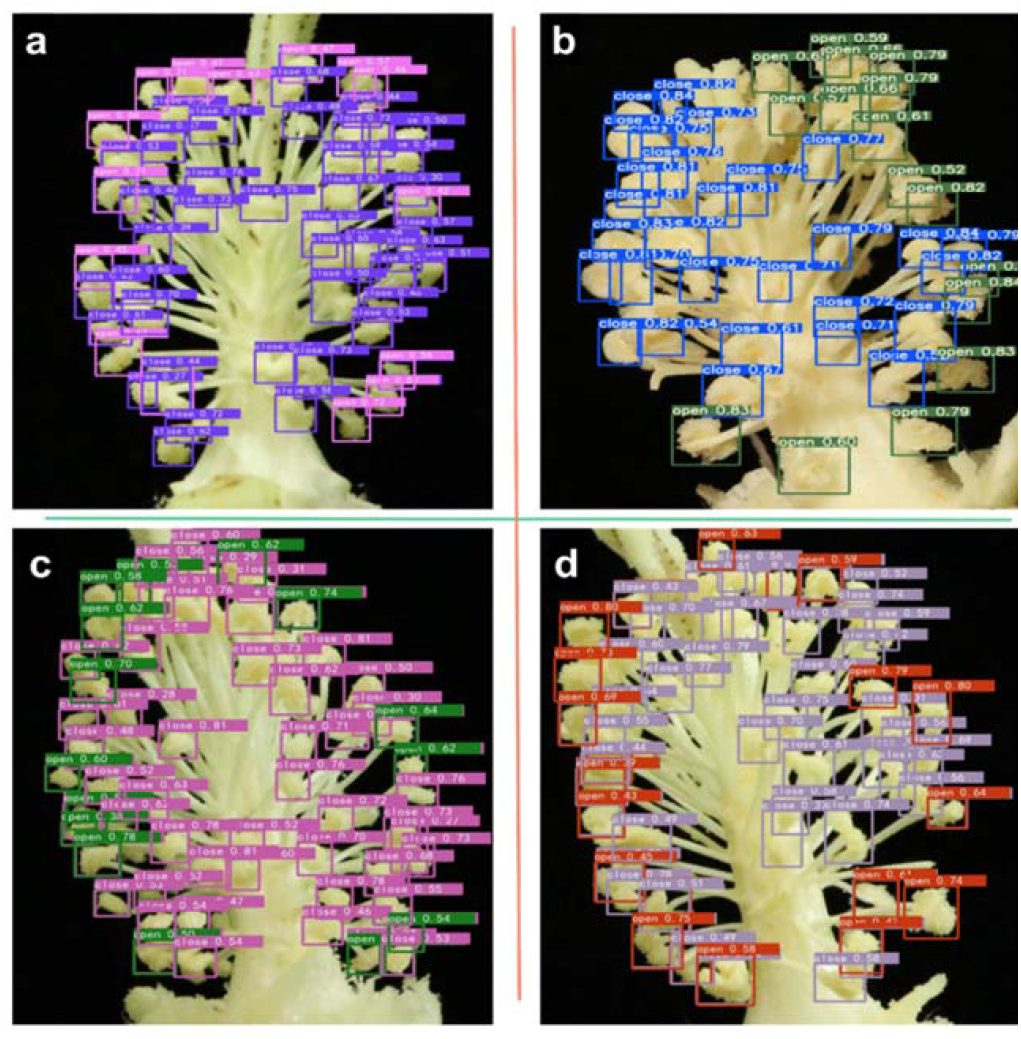
Cotton anther identification effect graph. **a**, The purple box marks an indehiscent cotton anther, and the pink box marks a dehiscent cotton anther. **b**, The blue box marks an indehiscent cotton anther, and the gray box marks a dehiscent cotton anther. **c**, The pink box marks an indehiscent cotton anther, and the green box marks a dehiscent cotton anther. **d**, The gray box marks an indehiscent cotton anther, and the red box marks a dehiscent cotton anther. In each test, the colors of the prediction boxes with different labels were randomly generated.

### Model comparison

#### Comparison of detection results of Faster R-CNN and YOLOv5

Faster R-CNN and YOLOv5 are used to train the same training set, the test results are compared on the same test set, and the correlation between the test results and the accurate value of manual labeling is analyzed. YOLOv5 using Darknet53 as the backbone network is a typical single-stage model, while Faster R-CNN using Res101 as the backbone network is a standard two-stage model. Obviously, YOLOv5 is more advantageous in detection speed. A comparison of the two models was shown in Figure 4a. Through training and validation, we found that mAP@0.5:0.95 of YOLOv5 was 0.485, while mAP@0.5:0.95 of Faster R-CNN was 0.478. In mAP@0.5:0.95, YOLOv5 was 0.007 higher than Faster R-CNN. In terms of the evaluation index of R^2^ in the validation set, the Faster R-CNN was 0.8712 in the category of “open” and 0.8373 in the category of “close”, and 0.82 in the category of “all”, which were 0.2523, 0.2619, and 0.3104 higher than YOLOv5, respectively. This may be due to the interference of location information. Although YOLOv5 has a slightly higher mAP@0.5:0.95, R^2^ is far lower than Faster R-CNN (Table S1). Since quantitative accuracy is our primary research goal, we decided to further optimize the two-stage model Faster R-CNN.

**Figure 4:**
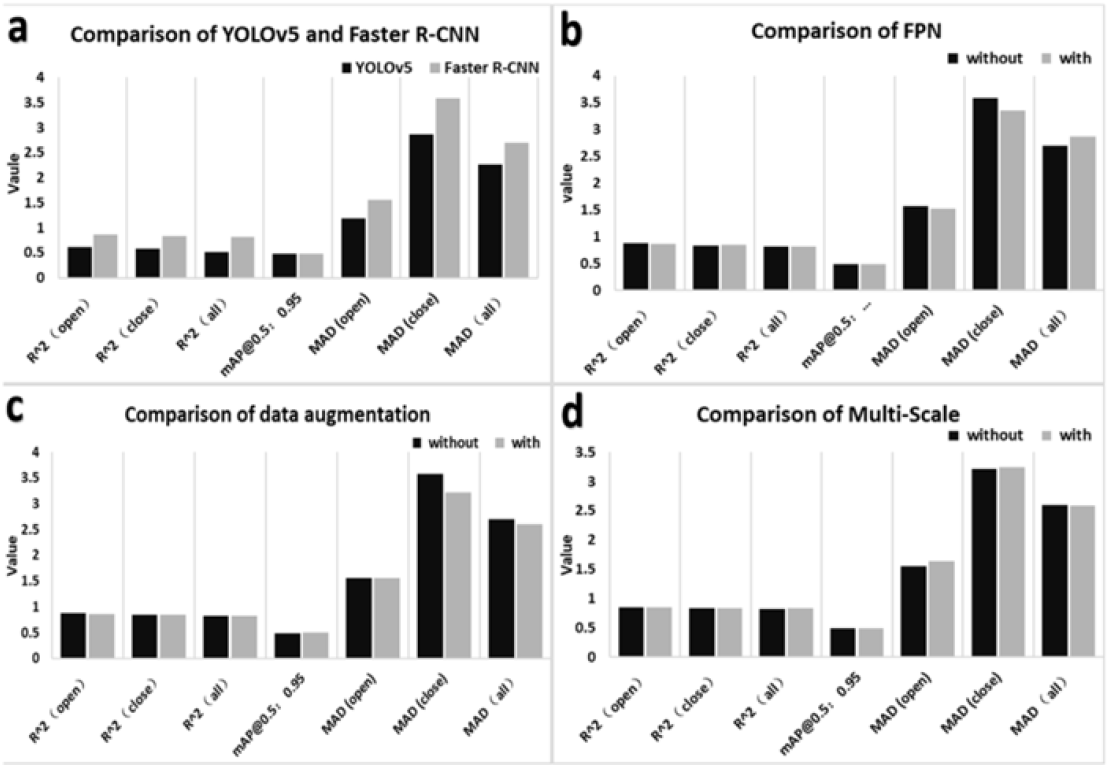
Comparison of different models. **a**, Comparison of YOLOv5 and Faster R-CNN. YOLOv5 model has higher recognition speed, Faster R-CNN model has higher detection accuracy. **b**, Comparison of with or without FPN (Feature Pyramid Networks) The mAP@0.5:0.95 of the improved model increased by 0.002, *R2* of “close” class increased by 0.003, and *R2* of “open” class and “all” decreased slightly. **c**, Comparison of with or without data augmentation. The improved model has a slight decline in the number of R2 in the open category and an improvement in other evaluation indicators. **d**, Comparison of with or without data Multi-Scale The result showed the mAP@0.5:0.95 of model was improved by 0.003 after Multi-Scale training. *R2* in the “open” and “close” categories fell by 0.0092 and 0.0007, respectively. *R2* in the “all” category went up 0.0086. “open” and “close” represent dehiscent and indehiscent anther, respectively.

#### Comparison of detection results with or without FPN (Feature Pyramid Networks)

To further improve the detection effect of the Faster R-CNN model, the FPN structure was added into the Faster R-CNN model. A comparison of the two models is shown in Figure 4b. The mAP@0.5:0.95 of Faster R-CNN with data augmentation was 0.48. For the R^2^ of the correlation of test value with the real value, Faster R-CNN with FPN structure have 0.8676, 0.8403 and 0.812 in the category of “open”, “close”, and “all”. Compared with that without FPN structure, mAP@0.5:0.95 of the improved model increased by 0.002 (Figure 5, model 1 and 3), R^2^ of “close” class increased by 0.003, and R^2^ of “open” class and “all” decreased slightly (Table S2).

**Figure 5:**
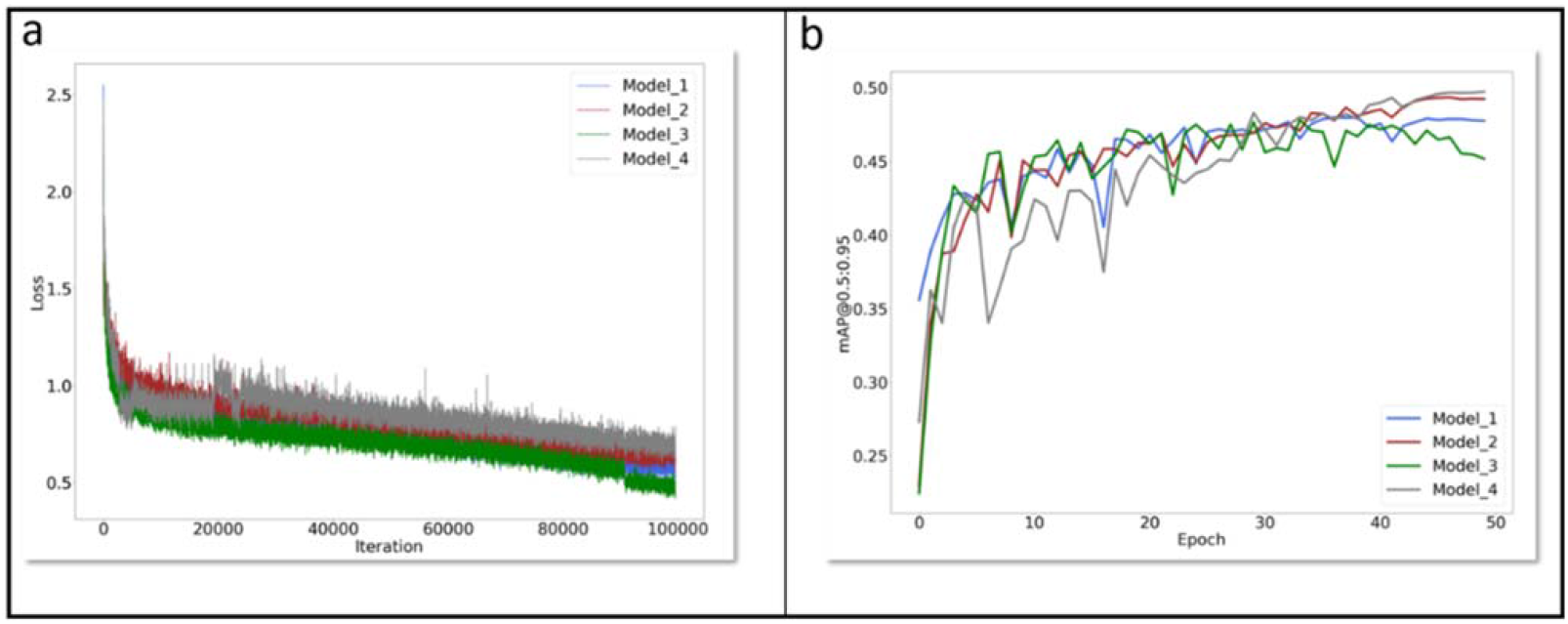
mAP@0.5:0.95 curves and LOSS curves. **Model 1** is the Faster R-CNN with FPN structure. **Model 2** is the Faster R-CNN with data augmentation and FPN structure. **Model 3** is the traditional Faster R-CNN. **Model 4** is the Faster R-CNN with Multi-Scale and data augmentation and FPN structure. **Epoch:** All the data were sent into the network to complete a process of forward calculation and back propagation. **mAP@0.5:0.95** is the process of increasing IoU from 0.5 to 0.95 according to the span of 0.05. The mAP corresponding to each IoU was added to obtain the average value of mAP in this process.

#### Comparison of detection results with or without data augmentation

The traditional Faster R-CNN model was taken without data augmentation. To avoid the effect of the sample imbalance, many kinds of data augmentation methods were added to the basic model, such as mixup, cutmix. The model with and without data augmentation were trained and tested on the same data set, and these detection results and correlations with the real value of manual labeling were compared. A comparison of the two models is shown in Figure 4c. We found that the mAP@0.5:0.95 of Faster R-CNN with data augmentation was 0.494, which was 0.016 higher than that of Faster R-CNN without data augmentation (Figure 5, model 1 and 2). For the R^2^ of the correlation of test value with the real value, Faster R-CNN with data augmentation were 0.8579, 0.8401 and 0.8235 in the category of “open”, “close”, and “all”, respectively. The R^2^ in the category of “close” and “all” of Faster R-CNN with data augmentation were 0.0028 and 0.0035 higher than that of Faster R-CNN without augmentation. While R^2^ in the “open” category of Faster R-CNN with data augmentation was 0.0133 lower than that of Faster R-CNN without data augmentation. Overall, the evaluation showed that the performance of Faster R-CNN with data augmentation is higher than that of Faster R-CNN without data augmentation (Table S3).

#### Comparison of detection results with or without Multi-Scale

To test whether the multi-scale training can improve the accuracy of the quantity of dehiscent anther, we added Multi-Scale on the basis of the traditional Faster R-CNN model. The specific content is to obtain the image pyramid at different scales, and then extract the features of different scales for each layer of images to obtain the feature map. Finally, the features of each scale are individually predicted. A comparison of the two models was shown in Figure 4d. The result showed the mAP@0.5:0.95 of model was improved by 0.003 after Multi-Scale training (Figure 5, model 4 and 2). R^2^ in the “open” and “close” categories fell by 0.0092 and 0.0007, respectively. R^2^ in the “all” category went up 0.0086. Thus, Multi-Scale training has a certain effect on our research goal of cotton anther identification (Table S4).

In this study, the change curves of each model in mAP@0.5:0.95 during the training process were shown in Figure 5. The peak value of traditional Faster-CNN mAP@0.5:0.95 curve was the lowest, while the peak value of Faster R-CNN model with data augmentation, Multi-Scale training and FPN structure was the highest. The loss curve of each model during the training process was shown in Figure 5. At the end of the training, the loss curve of the four models has tended to be stable.

#### Screening of HT tolerant cotton germplasms based on cotton anther phenotype data obtained using integrated Faster R-CNN model

In order to select high temperature (HT) tolerant cotton germplasms, the anther pictures of different cotton lines were obtained under normal temperature (NT) and HT. Then we counted the dehiscence status of anthers from 30 different cotton lines by manual observation and machine recognition, and the statistical results were shown in Table 1. The results of manual observation showed that the average dehiscence rate of cotton anthers treated with NT and HT were 84.35% and 35.46% respectively, and the results of machine recognition showed that the average dehiscent rates of cotton anthers treated with NT and HT were 83.81% and 35.08% respectively. First of all, we can believe that for the acquisition of the phenotypic data of cotton anther dehiscence rate, the result of machine recognition has been extremely accurate, and the recognition speed is fast, which is not affected by artificial subjective factors, save manpower and material resources, there are obvious advantages compared with manual observation. Secondly, there is a great difference in the anther dehiscent rate of the same cotton variety between HT and NT conditions. The result showed that HT greatly reduced the cotton anther dehiscent rate (Table 1), and then affected the pollination process, resulting in a reduction in cotton yield. Finally, by observing 30 cotton lines, we found that the anther dehiscent rate of S003 and S004 was still more than 85% under HT stress, which was significantly improved compared with other lines (Table 1). In addition, we screened cotton lines with HT tolerance in large quantities through machine recognition, and obtained more than 35 HT tolerant cotton lines. These HT tolerant germplasms will be used in cotton HT tolerance breeding.

**Table 1:** Screening of HT tolerant cotton germplasms using integrated Faster R-CNN model.

| Variety | Manual count |  |  |  |  |  | Machine count |  |  |  |  |  |
| --- | --- | --- | --- | --- | --- | --- | --- | --- | --- | --- | --- | --- |
|  | Normal temperature |  |  | High temperature |  |  | Normal temperature |  |  | High temperature |  |  |
|  | Open | Close | Dehiscent rate(%) | Open | Close | Dehiscent rate(%) | Open | Close | Dehiscent rate(%) | Open | Close | Dehiscent rate(%) |
| S001 | 39.5±2.02 | 5.75±0.94 | 87.17±2.3 | 34.75±4.21 | 15±4.76 | 70.41±8.26 | 38.25±2.39 | 5.5±1.32 | 87.31±2.97 | 35.25±2.92 | 13.75±3.9 | 72.47±6.85 |
| S002 | 26.25±3.49 | 17.25±1.37 | 78.6±2.25 | 9.25±2.14 | 17±2.2 | 65.68±3.41 | 26.75±3.75 | 7±1.29 | 79.35±2.25 | 17.25±1.31 | 8.5±1.76 | 68.06±3.13 |
| S003 | 42.25±2.92 | 11.25±2.01 | 78.84±3.82 | 30±6.17 | 4±1.58 | 86.55±5.03 | 40.25±2.86 | 11±1.87 | 78.54±3.25 | 30.25±5.76 | 5±1.82 | 84.46±5.56 |
| S004 | 32.25±2.28 | 3±1.22 | 91.86±3.02 | 27.25±0.62 | 2±0.91 | 93.27±3.02 | 32.25±2.28 | 2.25±0.75 | 93.63±1.92 | 27±0.7 | 2.25±0.75 | 92.4±2.44 |
| S005 | 27.25±3.9 | 9.75±3.94 | 77.32±7.54 | 29.25±2.81 | 18.25±5.12 | 36.19±5.06 | 26.75±4.42 | 10.25±3.59 | 74.14±6.48 | 17.5±4.13 | 29.25±2.95 | 36.12±4.43 |
| S006 | 28.75±1.38 | 6±1.08 | 83.97±2.31 | 18.25±1.37 | 13±4.22 | 61.04±9.65 | 28.25±1.44 | 5.5±0.65 | 83.86±1.01 | 18.5±0.86 | 12.75±3.9 | 61.75±8.76 |
| S007 | 20.75±3.59 | 5.75±1.75 | 76.92±8.33 | 16.25±2.53 | 11.5±1.19 | 42.3±4.65 | 19.75±2.68 | 6±1.68 | 76.08±7.37 | 10.75±0.63 | 18±2.27 | 37.94±1.81 |
| S008 | 17±3.08 | 6±0 | 72.69±3.06 | 12.25±0.75 | 18.5±1.19 | 39.95±2.81 | 18.25±3.25 | 5.75±0.25 | 74.95±2.57 | 10.5±0.28 | 19.5±0.64 | 35.03±0.95 |
| S009 | 25±2.85 | 4.5±1.7 | 86.35±3.53 | 13.5±0.5 | 17±2.94 | 45.62±5.17 | 22.5±2.72 | 5.5±1.5 | 80.64±4.45 | 13.25±0.75 | 15.5±2.25 | 46.97±4.96 |
| S010 | 24.25±2.56 | 5.5±1.44 | 82.31±3.49 | 12.5±1.44 | 21.75±2.17 | 36.39±1.82 | 23.75±2.75 | 6±1.58 | 80.87±3.52 | 11±0.57 | 21.5±1.93 | 34.04±1.27 |
| S011 | 24.25±4.49 | 4±0.91 | 85.81±1.41 | 8±2.85 | 24.5±6.73 | 19.77±7.44 | 23.5±4.34 | 3.25±0.62 | 87.28±1.94 | 8±2.67 | 23.25±6.14 | 20.43±7.04 |
| S012 | 28.25±2.52 | 2.5±0.5 | 91.51±1.97 | 0±0 | 29.25±8.6 | 0±0 | 26.5±1.84 | 2.25±0.62 | 91.9±2.57 | 0.5±0.5 | 28±8.33 | 4.545±4.54 |
| S013 | 27.5±6.06 | 3.75±0.75 | 86.79±3.53 | 0.25±0.25 | 45±5.11 | 0.5±0.5 | 24.75±5.45 | 4.25±0.47 | 84±3.15 | 0±0 | 43.75±3.19 | 0±0 |
| S014 | 29.75±4.53 | 6.75±2.09 | 81.08±5.81 | 3.25±3.25 | 37.25±1.88 | 7.22±7.22 | 28.5±3.79 | 6.5±2.21 | 81.08±6.17 | 3.75±3.42 | 38±2.44 | 8.164±7.44 |
| S015 | 32.25±1.03 | 3.75±1.1 | 90.01±2.61 | 17.25±4.26 | 20.75±3.01 | 44.65±8.45 | 31.5±0.64 | 3.75±1.1 | 89.76±2.71 | 16.75±3.49 | 18.75±1.43 | 45.98±5.93 |
| S016 | 22.75±3.47 | 4±1.58 | 86.06±6.08 | 14.75±0.63 | 7.25±1.03 | 67.39±3.2 | 22.25±3.19 | 4.5±1.19 | 83.23±4.37 | 14±0.58 | 7.5±1.04 | 65.48±3.34 |
| S017 | 32.75±2.39 | 8.25±4.44 | 81.68±9.2 | 12.5±1.04 | 22.25±4.39 | 37.12±3.56 | 32±2.27 | 8.5±4.42 | 80.75±9.66 | 12.5±1.19 | 22.25±3.68 | 36.64±2.97 |
| S018 | 31.5±1.5 | 2.25±0.47 | 93.42±1.31 | 10.75±0.95 | 21±2.35 | 34.07±1.61 | 30.5±0.95 | 2.75±0.47 | 91.75±1.32 | 12.75±0.75 | 21±1.08 | 37.82±2.19 |
| S019 | 22.75±1.03 | 3.25±1.18 | 88.39±4.22 | 14.25±1.49 | 10±1.63 | 58.91±5.47 | 22±0.91 | 3.5±1.32 | 87.41±4.48 | 13.75±1.11 | 10.5±1.85 | 57.4±4.71 |
| S020 | 29±3.1 | 4.25±1.18 | 86.98±3.78 | 0.75±0.48 | 29.5±8.53 | 3.55±2.8 | 26.25±1.93 | 3.75±1.03 | 87.58±3.06 | 0.5±0.5 | 30±6.94 | 2.63±2.63 |
| S021 | 28±0.41 | 3±1.08 | 90.58±3.16 | 10.75±3.09 | 22.75±6.34 | 32.5±6.02 | 29.57±0.85 | 3±0.91 | 90.96±2.6 | 9.75±2.93 | 23±5.4 | 29.34±4.63 |
| S022 | 29.5±5.1 | 2±0.91 | 93.95±2.75 | 0.25±0.25 | 45.25±3.09 | 0.51±0.51 | 28±4.45 | 1.75±1.03 | 94.65±3.17 | 0±0 | 43.75±3.19 | 0±0 |
| S023 | 26.25±2.09 | 2.5±0.28 | 91.06±1.51 | 0.5±0.28 | 43.25±6.42 | 1.045±0.6 | 25.25±1.37 | 2.5±0.28 | 90.91±1.19 | 0.5±0.28 | 43.5±6.33 | 0.994±0.57 |
| S024 | 30±4.65 | 6.75±1.7 | 80.44±5.82 | 0.75±0.48 | 37.75±2.66 | 2.07±1.43 | 28.5±3.79 | 6.5±2.21 | 81.08±6.17 | 0.75±0.75 | 38.5±3.88 | 2.34±2.34 |
| S025 | 18.75±3.32 | 7±0.7 | 71.34±5.16 | 13.5±0.65 | 21.75±3.33 | 39.36±4.57 | 19.25±3.17 | 8.5±1.19 | 68.31±5.9 | 14.5±0.29 | 22.75±3.17 | 39.82±3.47 |
| S026 | 32.5±1.04 | 3.75±1.1 | 89.99±2.76 | 3±1.68 | 41.75±7.9 | 9.35±6.76 | 31.5±0.64 | 3.75±1.1 | 89.76±2.71 | 3±2.34 | 42.25±8.73 | 9.669±8.28 |
| S027 | 23±3.82 | 4.75±1.37 | 83.29±3.76 | 13.5±1.19 | 7.25±0.75 | 65.03±2.37 | 22.25±3.19 | 4.5±1.19 | 83.23±4.37 | 14±0.58 | 7.5±1.04 | 65.48±3.44 |
| S028 | 31.25±2.89 | 4.25±1.49 | 87.53±4.6 | 5.25±1.93 | 20.5±3.96 | 19.19±4.81 | 31±1.87 | 4.75±1.18 | 86.58±3.46 | 5.25±2.75 | 21.25±4.47 | 14.49±7.04 |
| S029 | 9.25±3.07 | 2.75±0.25 | 72.05±6.26 | 0.5±0.5 | 10±0.82 | 3.57±3.57 | 8.5±3.18 | 2.5±0.29 | 71.18±6.45 | 0.5±0.29 | 10±0.91 | 4.42±2.6 |
| S030 | 21.75±2.8 | 4.25±0.25 | 83.15±1.67 | 11.25±0.25 | 18.25±0.48 | 38.15±0.57 | 20.75±1.93 | 4±0.4 | 83.79±1.07 | 11.25±0.85 | 17±1.35 | 39.87±0.67 |
| Average | 27.2±0.36 | 4.95±0.55 | 84.35±1.23 | 11.21±0.77 | 22.03±0.96 | 35.46±1.61 | 26.25±0.31 | 4.98±0.5 | 83.81±1.17 | 11.11±0.68 | 21.97±0.81 | 35.08±1.76 |

Open: the average number of dehiscent anthers from different flowers, n>4; Close: the average number of indehiscent anthers from different flowers, n>4; Dehiscent rate: the number of dehiscent anthers of each flower/(the number of dehiscent anthers of each flower + the number of indehiscent anthers of each flower), n>4.

## Discussion

Through analysis, we found that the mAP@0.5:0.95 value of the model increased significantly after adding data augmentation and FPN structure and Mulit-Scale, but the change of R^2^ was not significantly positively correlated with mAP@0.5:0.95. In order to obtain the most accurate data in the application, four models were trained, as shown in Figure 5 and tested them on the same batch of test sets. The recognition results obtained were integrated by the following formula:

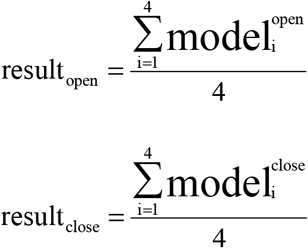

Among those, i represents the number of the model in Figure 5. Model_i_^open^ represents the number of dehiscent cotton anthers identified by modeli in the verification set. Model_i_^close^ represents the number of indehiscent cotton anthers identified by model_i_ in the verification set.

After the comparison with the real value, it is found that when the model is integrated, the detection result after ensemble effectively compensates for the error, and the correlation between the detection result and the real value will increase. After the ensemble of the four models, R^2^ of “open” reaches 0.8765, R^2^ of “close” reaches 0.8539, R^2^ of “all” reaches 0.8481, higher than the prediction result of either model alone. Therefore, when accurate data is needed, we can choose to integrate the detection results of the four models, so that the detection data obtained is the most reliable. Of course, directly using Faster R-CNN model with FPN structure and data augmentation and multi-scale has higher robustness and higher accuracy.

It is well known that anther is the male organ of plant, anther abortion will directly lead to male sterility and reduce yield. Our previous studies could be preliminarily concluded that HT stress can reduce cotton yield by inhibiting cotton male fertility. HT mainly decreased pollen viability, the anther growth number, and the percentage of dehiscent anther, caused the decreases of male fertility in cotton (Min et al., 2014; Ma et al., 2018). So far, no genes involved in HT affecting cotton male fertility have been cloned. Thus, further molecular biological methods can respond to this mechanism from the perspective of genes and improve cotton crop yield. Through the trained augmentation Faster R-CNN rapid identification system of cotton anther phenotype, can quickly investigate the anther phenotype and used to location of the genes affecting cotton anther dehiscence under HT. This will effectively promote HT resistance breeding of cotton and ensure cotton safe production under the trend of global warming.

### Conclusions and future directions

#### Conclusions

1. A cotton anther phenotype recognition system based on deep learning is proposed for the first time, which can help researchers to quickly investigate the anther phenotype of cotton and locate the genes that respond to the influence of stress on cotton anther for breeding improvement.
2. A lightweight cotton anther dehiscence detection model based on YOLOv5 is proposed, which can be easily implanted into embedded devices or mobile devices.
3. Through the change of the accuracy and correlation of Faster R-CNN after the improvement of the data augmentation method, the feasibility and superiority of the improved method were verified. Model enhanced by data.
4. After the ensemble of the four models, R^2^ of “open” reaches 0.8765, R^2^ of “close” reaches 0.8539, R^2^ of “all” reaches 0.8481, higher than the prediction result of either model alone, and can completely replace the manual counting method. It provides new technical support for cotton reproductive development and HT tolerance breeding.

#### Future directions

In this study, YOLOv5 and Faster R-CNN were applied to identify the dehiscence state of cotton anther, and achieved fast and accurate identification. But there are still some areas where there is room for improvement:

1. We only examined the dehiscence of cotton anthers, but other phenotypes such as the growth position of anthers and the distance between anthers and stigmas are also important for the reproductive development of cotton. Other phenotypic characteristics of cotton anther can be collected by using a comprehensive platform that integrates multiple data points to analyze cotton reproductive development.
2. The cotton anther dehiscence recognition model trained in this study should be further developed and applied to other computer platforms or servers to facilitate cotton reproductive development researchers to use the model to quickly obtain anther dehiscence data.
3. In this study, the experience of deep learning model training for cotton anther dehiscence can be applied to other plant anther state detection. It is one of the directions to further enrich the research content to further construct cotton anther state recognition models of various crops based on deep learning.

## Materials and methods

### Material growing and dataset acquisition

In total, 510 cotton lines from natural populations were planted in 2016-2019 in experimental cotton fields at Huazhong Agricultural University, Wuhan, Hubei (113.41E, 29.58N), Turpan, Xinjiang (89.19E, 42.91N), and Alar, Xinjiang (81.29E, 40.54N). At Wuhan, the field was planted at a density of 27,000 plants per hectare with each row including more than 12 individuals. At Alar and Turpan, Xinjiang, the fields were set up with two streets and planted at a density of 195,000 plants per hectare. More than 30 individuals of each line were arranged in rows. Cotton anther images were collected each year at each location three days after the onset of normal temperatures and after high temperatures during bloom.

A Canon 70 d HD digital camera was used throughout the acquisition of a research image set. To prevent the negative interference of background with the subsequent machine recognition effort, a black curtain was used as the photo background for this experiment. In the actual image collection process, it was found that the cotton anthers were surrounded by cotton petals, and the anthers growing at the root of the style were not easily captured by the camera, so it was not conducive to the accurate collection of data to take the pictures directly. Thus, it was necessary to preprocess the cotton flowers before acquiring the pictures by stripping the cotton petals and fixing the anther sides. To prevent overfitting, the model due to insufficient training data, the same anthers were included in multiple distant near-field images (Figure 6). Finally, a total of 2,845 high-definition RGB anther images were acquired.

**Figure 6:**
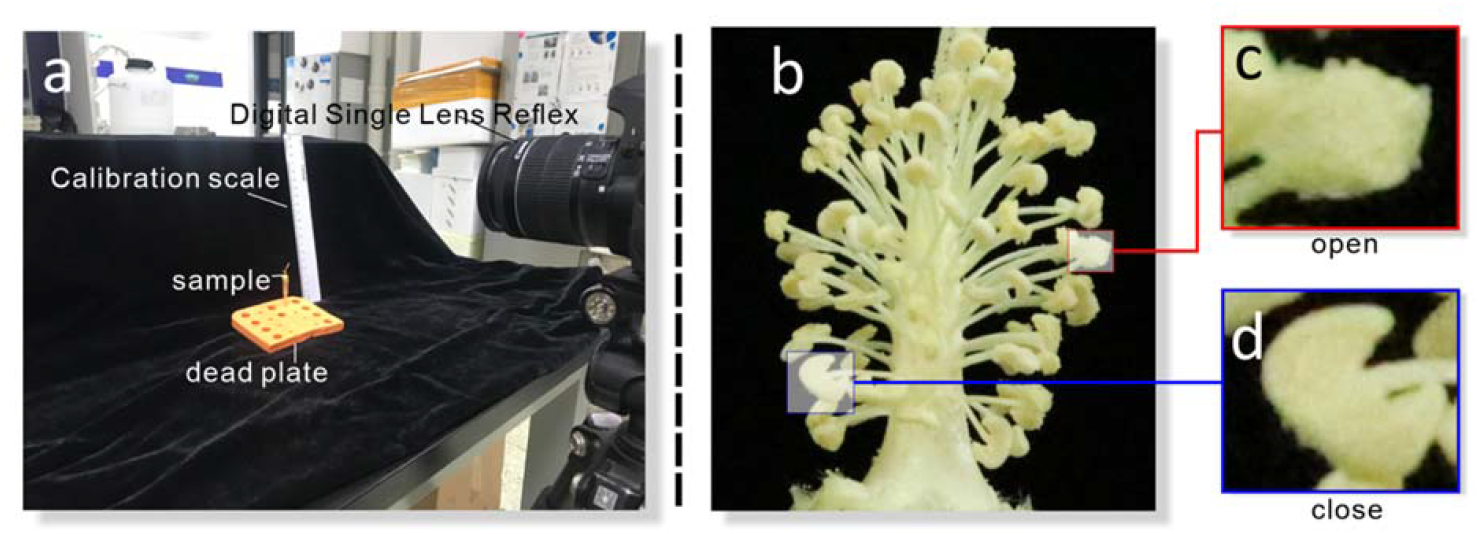
Data acquisition. **a:** The image dataset captures the platform scene. **b:** Image of cotton anther. **c:** The surface of dehiscent cotton anther (open) is rough from the image. **d:** The surface of indehiscent cotton anther (close) is smooth from the image.

Morphologically, dehiscent anthers are rough and grainy because pollen is released and adheres to anther edges, while indehiscent anthers have smooth edges, because no pollen is released. Therefore, the obtained cotton anther images were annotated using Labeling image annotation software, as shown in Figure S1. The image boundary of each visible cotton anther is within an annotation box that reduces the influence of background on model training and is labeled “open” and “close” to distinguish dehiscent and indehiscent anthers, respectively. A total of 2,845 images were annotated one by one. The images were used as the data set and were randomly divided into a training set and validation set in the ratio of 7:3 (Table S5).

### Experimental operation environment

The hardware environment used in this study is in Table S6, and on this basis, the training environment is python, open-cv, cuda, etc., and the frameworks used in this study are paddle and pytorch.

### Metrics used to evaluate the proposed method

In this study, we use mAP@0.5:0.95, as well as MAD (Mean Absolute Deviation) and R^2^ as the evaluation indicators of the model. The indicators are explained as follows:

mAP@0.5:0.95 is the process of increasing IoU from 0.5 to 0.95 according to the span of 0.05. The mAP corresponding to each IoU is added to obtain the average value of mAP in this process. The formula is expressed as follows:

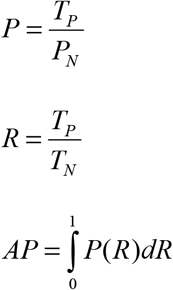

In the above formula, T_P_ is the correct number of a category identified by the model, P_N_ is the total number of categories identified by the model, and T_N_ is the true number of a category. Averaging the AP values of all categories is called mAP.

We took the absolute value of the absolute error between the measured value and the real value and then calculate the average value, and called it the MAD. Due to the deviation is absolute value, the positive and negative will not be offset, thus the mean absolute error can reflect the actual situation of predicted value deviation. The smaller the value, the closer the prediction of model is to reality.

The main purpose of this study is to develop a deep learning model that can quickly and accurately identify anther dehiscence and explore the influence of high temperature stress on cotton anther dehiscence. In the model identification phase, we identify the location of the cotton anther without strict requirements, but we demand model to recognize the number with the result of artificial observation is the most close to, so the number by artificial observation to anther as accurate value of validation set was used, with correlation coefficient between predicted values and the accurate value as the main evaluation index of the model.

## Supplemental Materials

**Figure S1: Image labeling**

The obtained cotton anther images were annotated using “Labeling” image annotation software. Green boxes represent indehiscent anthers and red boxes represent dehiscent anthers. When the image labeling was finished, we corresponded the location information of the image with the name of the image one by one and saved it in VOC format.

**Table S1:** Comparison of YOLOv5 and Faster R-CNN

**Table S2:** Comparison of FPN

**Table S3:** Comparison of data augmentation

**Table S4:** Comparison of Multi-Scale

**Table S5:** Dataset

**Table S6:** Experimental configuration

## Acknowledgements

Appreciations are given to the editors and reviewer.

